# Identification of a cytotoxic peptide from the spider *Enoploctenus cyclothorax* venom on leukemia cell lineages

**DOI:** 10.64898/2025.12.27.696694

**Authors:** Clara Sanches Bueno, Gabriela Borges Vieira, Juliana Mozer Sciani, Thomaz Augusto Alves da Rocha e Silva

## Abstract

**Background:** Chronic myeloid leukemia and acute lymphoblastic leukemia are significant malignancies requiring novel therapeutic agents. This study aimed to identify cytotoxic components of the venom of the spider *Enoploctenus cyclothorax* against K562 and NALM-6 cell lines.

**Methods:** Venom was extracted, lyophilizated and incubated with cultured K562 and NALM-6 cells at concentrations ranging from 0.1 to 100 µg/mL for 24 and 48 hours. MTT assay for cell viability was assessed to measure cytotoxicity. Venom fractionation started with centrifugation on AMICON® filters in a cutting edge of 10 kDa which separates low-(LW) and high-molecular weight (HW) fractions. The active fraction was fractionated using reversed-phase HPLC and the most active compound was purified and analyzed via mass spectrometry to determine its molecular weight.

**Results:** Cytotoxic activity was observed in both cell lines at concentrations of 10, 30, and 100 µg/mL *E. cyclothorax*. The centrifugation revealed that antineoplastic activity was concentrated in HW fraction. Subsequent chromatography isolated fraction F7 (specifically F7c) as the most potent. Mass spectrometry characterized F7c as a mild hydrophobic polypeptide with a molecular mass of approximately 6270 Da.

**Conclusion:** A polypeptide of approximately 6270 Da isolated from *E. cyclothorax* venom demonstrates significant *in vitro* cytotoxic properties against leukemic lineages. Further sequencing and synthesis are required to advance the pharmacological bioprospecting of this potential antineoplastic agent.

## 1. Introduction

Chronic Myeloid Leukemia (CML) and Acute Lymphoblastic Leukemia (ALL) represent significant challenges in modern oncology due to their high prevalence and economic burden. CML accounts for 15–20% of adult leukemias, with prevalence rates expected to rise dramatically by 2050 (Bortoelho et al., 2008; Höglund et al., 2015). Conversely, ALL is the leading cause of childhood malignancies, with over 111,000 prevalent cases estimated in the United States alone in 2020 (Pedrosa & Lins, 2002; SEER NIH, 2024). While current treatment protocols involving traditional chemotherapy and targeted therapies have improved survival, they are frequently limited by systemic toxicity, severe side effects, and the development of resistance (Kayser et al., 2021). Consequently, the identification of novel therapeutic agents remains a critical priority.

Arachnid venoms have emerged as a promising source of bioactive molecules with antineoplastic properties.^1^ Specifically, the Ctenidae subfamily produces venoms characterized by proteolytic, hyaluronidase, and phospholipase activities (Okamoto et al., 2009).^2^ Building upon recent findings by our group, which successfully purified a cytotoxic toxin from the Ctenidae family (de Mato et al., 2023), this study investigates the potential cytotoxic and cytostatic activity of venom from *Enoploctenus cyclothorax*.

## 2. Objectives

The primary objective of this study was to evaluate the cytotoxic and cytostatic potential of the venom from *Enoploctenus cyclothorax* against leukemia cell lineages, followed by the purification and identification of the specific toxins responsible for these activities.

## 3. Methods

### 3.1 Venom Extraction and Preparation

Enoploctenus cyclothorax specimens were collected at the Alto Ribeira Touristic State Park (PETAR, Brazil). Venom extraction was performed according previous described method (Rocha e Silva et al, 2009). Crude venoms were centrifuged at 3000 x g, lyophilized, stored at -80°C, and filtered (0.22 µm) prior to use. Animals were released into their natural habitat post-recovery.

### 3.2 Cell Culture

NALM-6 (Acute Lymphoblastic Leukemia, DSMZ no. ACC 128) and K562 (chronic myelogenous leukemia, ATCC CCL-243) cell lines were used as models. Cells were cultured in RPMI-1640 medium (Gibco®, NY, USA) supplemented with 10% fetal bovine serum and 1% antibiotic-antimycotic solution (Gibco®, NY, USA) at 37°C in a 5% CO_2_ atmosphere. Subculturing occurred every 72 hours. Prior to assays, viability was confirmed >95% using the Trypan blue exclusion method.

### 3.3 Cytotoxicity Assays

To assess antineoplastic activity, cell cultures were incubated with venom concentrations of 0 (control), 0.1, 0.3, 1, 3, 10, 30, and 100 µg/mL for periods of 24 and 48 hours. Cytotoxicity was quantified using the MTT (3-(4,5-dimethylthiazol-2-yl)-2,5-diphenyltetrazolium bromide) colorimetric assay. Absorbance was measured using a Varioskan spectrophotometer, and data were analyzed via polynomial regression to determine cell viability curves.

### 3.4 Venom Purification and Identification

Following the confirmation of bioactivity, venom fractionation was performed using a Shimadzu Prominence Pro chromatograph equipped with a Supelco C18 column. Elution used a linear gradient of 0–100% Buffer B (90% acetonitrile + 0.1% TFA) in Buffer A (water + 0.1% TFA) over 100 minutes. Active fractions were re-chromatographed using a 0–15% B gradient over 30 minutes. The molecular weight of the isolated toxin was determined inserting an aliquot of 5 μL in the mass spectrometry (Xevo GS QToF, Waters Co., Milford, MA, USA) by direct infusion, with elution of 50% acetonitrile and 0.1% formic acid, 0.2 mL/min, in positive ESI ionization mode. Data were collected in a full MS scan ranging from 100 to 1200 m/z, and manually analyzed.

### 3.8 Ethical and Legal Aspects

This study complied with Brazilian regulations (Federal Law No. 11.794/08). Spiders collection and access to venom were authorized under federal licenses SISBIO 58786-1, CGen 010115/2014-5, and state licence COTEC 260108–004.610/2016.

## 4. Results

### 4.1 Cytotoxicity Screening and Venom Selection

*Enoploctenus cyclothorax* crude venom exhibited dose-dependent antiproliferative activity against NALM-6. Significant effects were observed at concentrations of 10, 30, and 100 µg/mL. At 10 µg/mL, a reduction in the growth rate was noted (cytostatic effect), while concentrations of 30 and 100 µg/mL caused a reduction in cell numbers below the initial input, indicating cytotoxicity (Figure 1).

**Figure 1.**
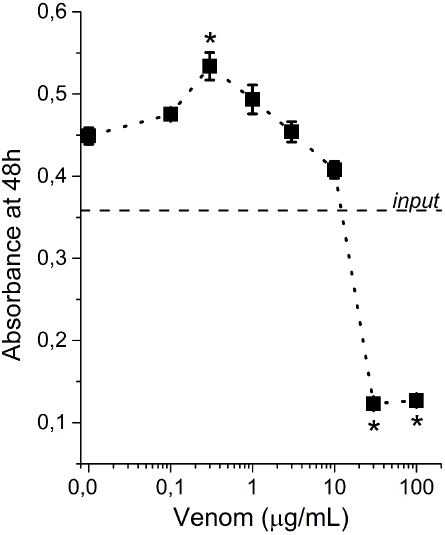
Cytotoxic activity of *Enoploctenus cyclothorax* against Nalm-6 cell lineage at 48h. At concentrations of 30 and 100 µg/mL; the graph demonstrates a drop in cell count to values below the initial input. *p<0,05 compared to 0,0 venom concentration.

### 4.2 Bioassay-Guided Fractionation

Separation of *E. cyclothorax* venom into light (LW < 3000 Da) and heavy (HW > 3000 Da) fractions revealed that antineoplastic activity was retained in the HW fraction, which exhibited a cytostatic profile similar to the crude venom. Cation exchange chromatography of the HW fraction yielded 16 peaks. Among these, Fraction 7 (F7) was the sole fraction displaying cytostatic activity against K-562 cells, maintaining cell numbers at baseline levels after 24 hours (Figure 2).

**Figure 2.**
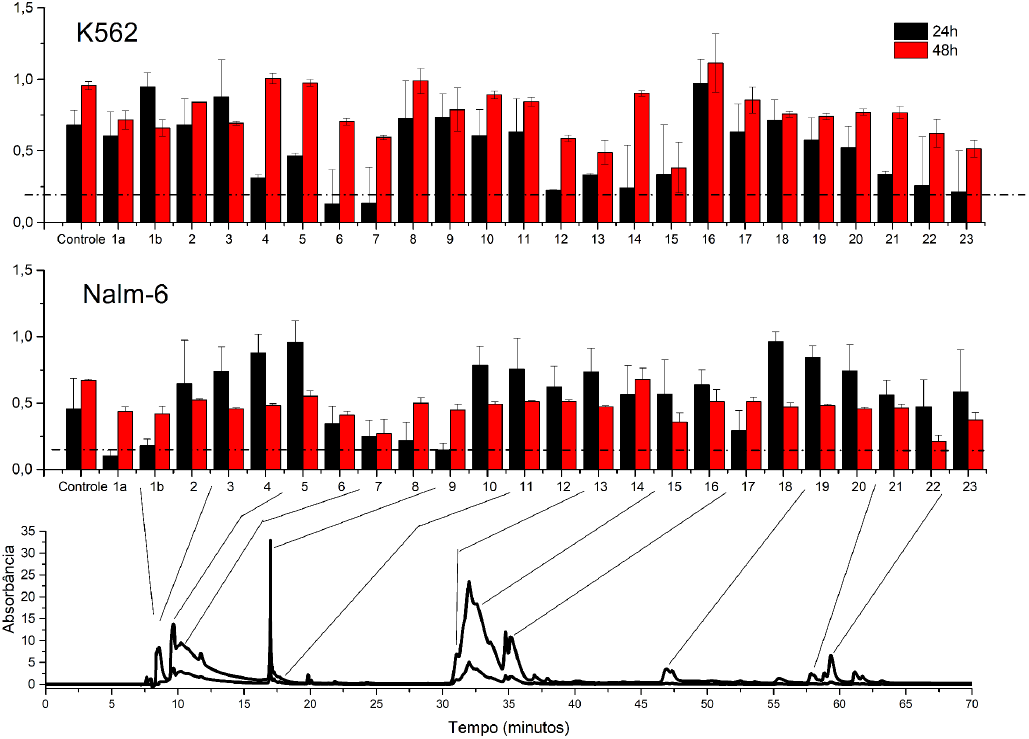
Cytotoxic profile chromatographic fractions of the venom of *E. cyclothorax*. Fraction 7 was elected for further characterization.

Subsequent reverse-phase chromatography of F7 isolated three sub-fractions (F7a, F7b, and F7c). Due to sample yield limitations, further testing was restricted to the NALM-6 cell line. Within this set, only sub-fraction F7c exhibited significant antineoplastic activity after 48 hours. (Figure 3).

**Figure 3.**
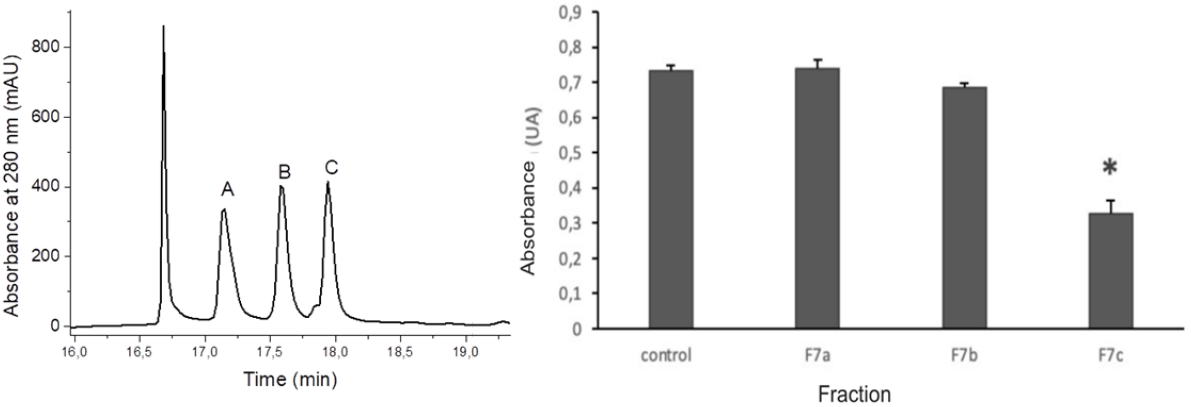
Reversed phased chromatography polishing of fraction 7 (left) demonstrating cytotoxic activity of subfraction 7c (right). *p<0,001.

Mass spectrometric characterization of the isolated bioactive peptide F7c determined a molecular weight of 6270 Da (Figure 4).

**Figure 4.**
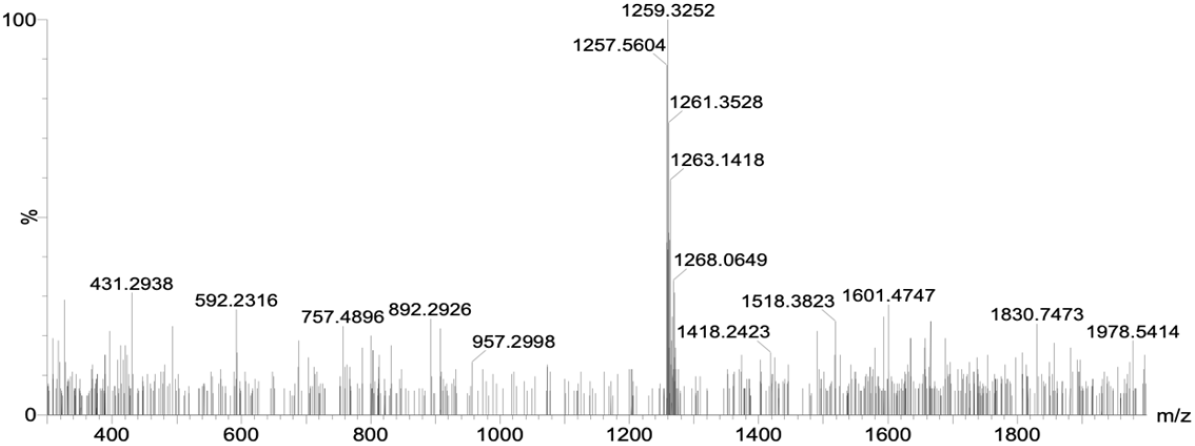
Section of the F7c fraction spectrum in Quadrupole-TOF-MS. A peak at 1259 m/z was detected and manually deconvoluted to a mass of 6270 Da.

## 5. Discussion

The unique ecological pressures exerted on cave-dwelling species, such as *Enoploctenus cyclothorax*, likely drive the selection of toxins with high specificity and potency (King, 2015). While previous studies have characterized proteolytic and phospholipase activities in the Ctenidae subfamily (Okamoto et al., 2009), this study represents the first specific characterization of their antineoplastic potential against leukemia cell lines.

Our results demonstrated that *E. cyclothorax* venom exhibits cytotoxic activity against human leukemia cells. Bioassay-guided fractionation localized the antineoplastic activity to the High Molecular Weight (HW) fraction (>3000 Da). This finding, corroborated by the isolation of the F7c toxin with a molecular mass of 6270 Da, suggests the active agent is likely a peptide or protein. This aligns with other arachnid-derived antineoplastic molecules, such as oxyopinins and phospholipase D, which target cell membranes or metabolic pathways (Heinen & Gorini, 2011; Belokoneva et al., 2003).

Notably, the potency of the isolated fraction highlights its therapeutic promise. The HW fraction of *E. cyclothorax* inhibited K-562 cell growth at a concentration of 1 µg/mL within 24 hours. In a comparative context, this concentration is five-fold lower than that required for *Macrothele raven* venom to achieve similar inhibition in the same cell line (Bedi et al., 1994; Liu et al., 2012). This superior potency suggests that *E. cyclothorax* venom may harbor novel molecules with optimized bioavailability or specificity profiles.

Despite the promising findings, this study presents limitations. First, the low yield of the purified sub-fraction F7c restricted the scope of the final bioassays; consequently, cytotoxic activity could only be verified in the NALM-6 cell line, precluding parallel validation in the K562 lineage. Second, while the molecular mass of the toxin was determined, its complete amino acid sequence remains to be elucidated, which is a prerequisite for peptide synthesis and detailed structure-activity relationship studies. Finally, the investigation was confined to *in vitro* leukemia models. The selectivity of the molecule against healthy cells was not assessed to rule out non-specific toxicity, nor were *in vivo* studies conducted yet to evaluate the compound’s bioavailability, pharmacokinetics, and systemic safety profile.

Although the yield of fraction F7c limited the extent of parallel testing, the identification of a 6270 Da toxin capable of significant cytotoxicity at low concentrations warrants further proteomic analysis and synthesis. These findings reinforce the potential of cave-dwelling arachnid venoms as a source of novel scaffolds for oncohematological drug development.

## 6. Conclusion

This study successfully isolated a bioactive fraction from *Enoploctenus cyclothorax* venom with a molecular weight of 6270 Da, indicative of a polypeptide structure. The toxin demonstrated significant cytotoxicity against leukemia cell lines, suggesting a potent mechanism of action. Further investigations will prioritize the full sequencing and synthesis of F7c, as well as structural modifications to optimize its pharmacological profile.

